# Awakening prime editing for precision engineering of probiotic *Escherichia coli* Nissle 1917

**DOI:** 10.1101/2024.10.28.620590

**Authors:** Pei-Ru Chen, Ying Wei, Xin Li, Hai-Yan Yu, Shu-Guang Wang, Xian-Zheng Yuan, Peng-Fei Xia

**Author notes:** Correspondence: Peng-Fei Xia, School of Environmental Science and Engineering, Shandong University, Qingdao 266237, China.

## Abstract

CRISPR-Cas systems are transforming precision medicine with engineered probiotics as next-generation diagnostics and therapeutics. To promote human health and treat disease, engineering probiotic bacteria demands maximal versatility to enable non-natural functionalities while minimizing undesired genomic interferences. Here, we present a streamlined prime editing approach tailored for probiotic *Escherichia coli* Nissle 1917 utilizing only essential genetic modules and an optimized workflow. This was realized by assembling a prime editor consisting of the CRISPR-Cas system from *Streptococcus pyogenes* with its native codons and a codon-optimized reverse transcriptase, and by orchestrating the induction levels. As a result, we achieved all types of prime editing in every individual round of experiments with efficiencies of 25.0%, 52.0% and 66.7% for DNA deletion, insertion, and substitution, respectively. A comprehensive evaluation of off-target effects revealed a significant reduction in unintended mutations, particularly in comparison to two different base editing methods. Leveraging the prime editing system, we developed a barcoding system for strain tracking and an antibiotic-resistance-gene-free platform to enable non-natural functionalities. Our prime editing strategy awakens back-to-basics CRISPR-Cas systems devoid of complex or extraneous designs, paving the way for future innovations in engineered probiotics.

## Introduction

CRISPR-Cas-based systems are revolutionizing gene editing across all three domains of life. However, the CRISPR-Cas gene editing with a “dead or alive” selection is challenging in bacteria, where the system was first discovered as an immune system against foreign DNAs. These challenges include the toxicity of Cas nucleases, the inefficient homology-directed repairing machinery, and the low capability of receiving foreign DNAs (1, 2). Despite inspiring advancements, these obstacles hinder further developments of genetic systems, especially in non-model or non-conventional bacteria, such as probiotics (3, 4), cyanobacteria (5), acetogens (6), and marine bacteria (7, 8), which often harbor unusual biological functionalities that benefit human health and sustainability. Innovative gene editing methodologies that overcome or bypass these biological barriers are imperative.

Prime editing and base editing are emerging CRISPR-Cas systems that allow precision genetic manipulation without generating double-strand breaks or requiring donor DNAs (9, 10). By deploying a “search-and-replace” concept (11), these methods can circumvent the aforementioned biological challenges and, theoretically, enable efficient gene editing in bacteria at a single nucleotide resolution (11-13). Base editing has been successfully applied to various bacteria with unique characteristics (e.g., *Synechococcus elongatus, Clostridium ljungdahlii, Streptomyces collinus*, and *Roseovarius nubinhibens*), where early STOP codons can be installed to inactivate genes of interest (GOIs) with nearly 100% efficiencies (6, 7, 14, 15). However, its editing capabilities are constrained by the intrinsic working mechanism of deamination, and off-target events become a trade-off for high editing efficiency (16-18). As one step forward, prime editing enables insertion, deletion, and replacement of small DNA fragments, and allows all twelve kinds of base changes with significantly reduced off-target events (19). These advantages make prime editing an ideal method for genetic manipulation and have been demonstrated in various cell types, including human, mammalian, and plant cells. The few reports (20-23), nevertheless, imply unexpected difficulties in developing prime editing systems in bacteria.

Engineered probiotics, as next-generation diagnostics and therapeutics, are expanding the horizon of clinical innovations by combining their natural health-promoting features and designed non-natural biological functionalities. *Escherichia coli* Nissle 1917 (EcN), a probiotic isolate from over 100 years ago, can colonize the gastrointestinal tract and treat conditions such as diarrhea and infectious inflammation (24). EcN has been engineered to sense and eliminate pathogens (25, 26), convert dietary compounds into anti-cancer agents (27), and deliver immunotherapeutics in situ for the treatment of colorectal neoplasia (28), showcasing great potential for medical applications. Like other isolated bacteria from nature, EcN is resistant to genetic manipulation, preventing further clinical progress (29). To fully uncover its potential, a customized gene editing system is needed to enable versatile and precision DNA manipulation in the genome with minimal on-target byproducts and off-target consequences.

Here, we present a modularly designed and optimized prime editing system for the engineering of EcN with only essential genetic modules. Efficient gene editing has been achieved and demonstrated by manipulating the amino acid metabolism. The off-target effects were evaluated and compared with base editing systems. By using prime editing, we created a barcoding system for the identification and quantification of engineered EcN, and we established an antibiotic-resistance-gene-free (ARG-free) platform to enable non-natural functionalities. Our study not only provides a tailored gene editing approach for probiotic bacteria, but also offers new insights into the development of streamlined CRISPR-Cas systems that overcome inherent biological barriers in bacteria.

## Results

### Modularly design of the prime editing system

A prime editing system consists of a prime editor, a fusion of Cas9 nickase (nCas9) and reverse transcriptase (RT), and a programmable prime editing guide RNA (pegRNA). The pegRNA contains a normal guide RNA (gRNA) for navigation, a primer binding site (PBS) and a reverse transcription template (RTT) containing the intended edits (11). To perform prime editing, the prime editor first finds the target DNA sequences and makes a nick (single-strand DNA break) **(Figure 1A)**. Then, the PBS hybridizes to the nicked DNA strand, and the reverse transcription of RTT is initiated to produce the donor DNA. After flap equilibration, cleavage and ligation processes, the designed edits will be incorporated into the genome **(Figure 1A and Figure S1)**. Accordingly, we designed a customized single-plasmid system with four modules and generated the working plasmid serial pRC **(Figure 1B and Table S1)**. Module 1 encodes the prime editor containing nCas9 from *Streptococcus pyogenes* and the engineered RT from Moloney murine leukemia virus under the control of a *lacI*-P_trc_ inducible system **(Figure 1B)**. Two different versions of DNA sequences of nCas9, one from the published PE2 (hereafter nCas9H) (11), and one of the original sequence from *S. pyogenes* (hereafter nCas9S), were employed, and RT was codon optimized for *E. coli* **(Table S2)**, generating two prime editors, PE.H (with nCas9H) and PE.S (with nCas9S), respectively. Module 2 consists of the pegRNA driven by a constitutive P_J23119_ promoter, and module 3 encodes a temperature-sensitive origin of replication for the controllable replication and curing of working plasmids. For module 4, we employed a gentamicin resistance gene for the selection and maintenance of plasmids.

**Figure 1.**
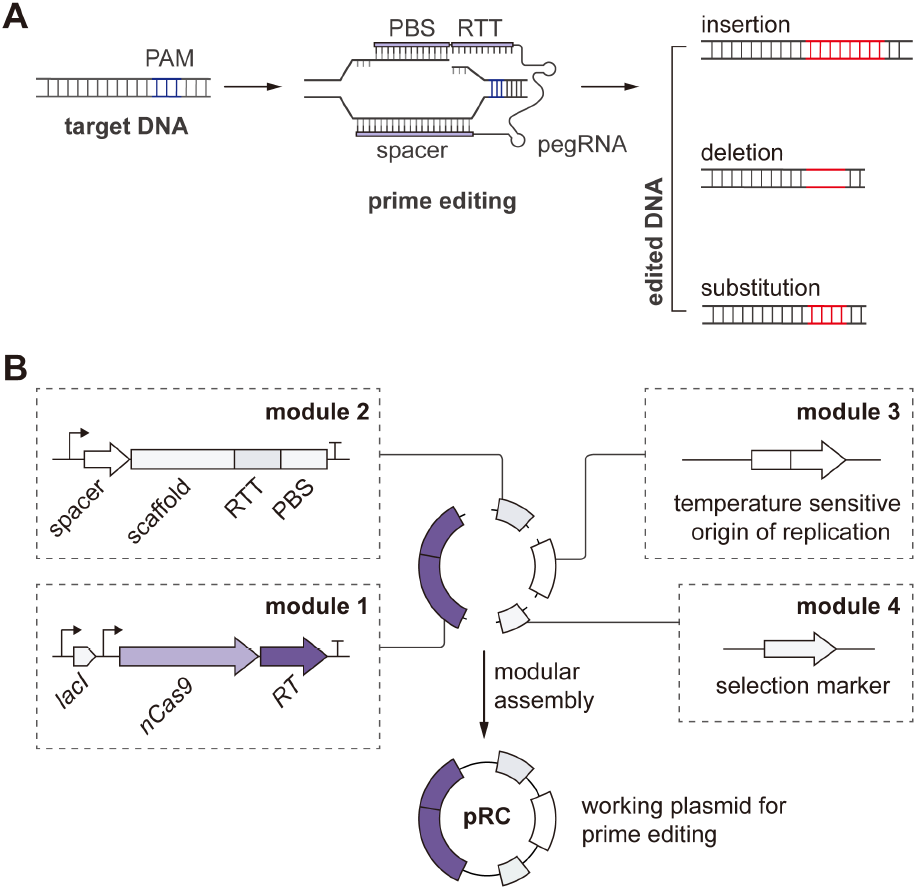
Prime editing system for EcN. **(A)** Schematic principle of prime editing. In brief, the prime editor introduces a nick on the target DNA and incorporates the designed edits with a specific pegRNA. **(B)** Modular design of the working plasmid serial pRC. For module 1, the nCas9 and RT are controlled by an inducible promoter *lacI*-P_*trc*_ driven by *lacI*. For module 2, the pegRNA is controlled by a constitutive promoter P_J23119_. The vector contains a temperature-sensitive origin of replication as module 3 and a selection marker as module 4. PBS, primer binding site; RTT, reverse transcription template; RT, reverse transcriptase.

### Extended induction enables prime editing in EcN

We chose *glnA*, encoding the glutamine synthetase, as a target and designed specific pegRNAs for DNA insertion (pegRNA01), deletion (pegRNA02) and substitution (pegRNA03) **(Figure 2A and Table S3)**. As a result, we successfully achieved DNA deletion with an efficiency of 11.3 ± 12.7% with pRC02 (PE.H with pegRNA02) and 4.0 ± 4.6% with pRC05 (PE.S with pegRNA02) **(Figure 2B, 2C, and Table S1)**, whereas we could not obtain the designed editing in every round of experiments. No designed insertion or substitution of nucleotides was identified **(Figure 2B)**. We hypothesized that the failed editing was due to the inadequate expression of the prime editors. Then, we extended the induction time from 24 h to 48 h, and successful editing with PE.H was achieved with increased efficiencies for insertion (10.4 ± 7.2%), deletion (13.1 ± 12.5%) and substitution (11.0 ± 19.2%) **(Figure 2B)**. However, DNA deletion and substitution could still not be obtained in each experiment. We observed better editing performances of PE.S. The editing efficiency of deletion increased to 25.0 ± 17.7%, which is 6.25-fold higher (P = 0.076) than that of 24 h induction **(Figure 2C)**. The editing efficiencies of insertion and substitution reached 52.0 ± 13.0% and 66.7 ± 13.0%, respectively, which were significantly higher than those induced for 24 h **(Figure 2C)**. These edits would lead to the inactivation of *glnA* by shifting the ORF or inserting early STOP codons **(Figure 2D)**. As expected, all the edited EcN exhibited the desired phenotypes that they could not grow in the M9 minimal medium without the supplement of glutamine **(Figure 2E)**.

**Figure 2.**
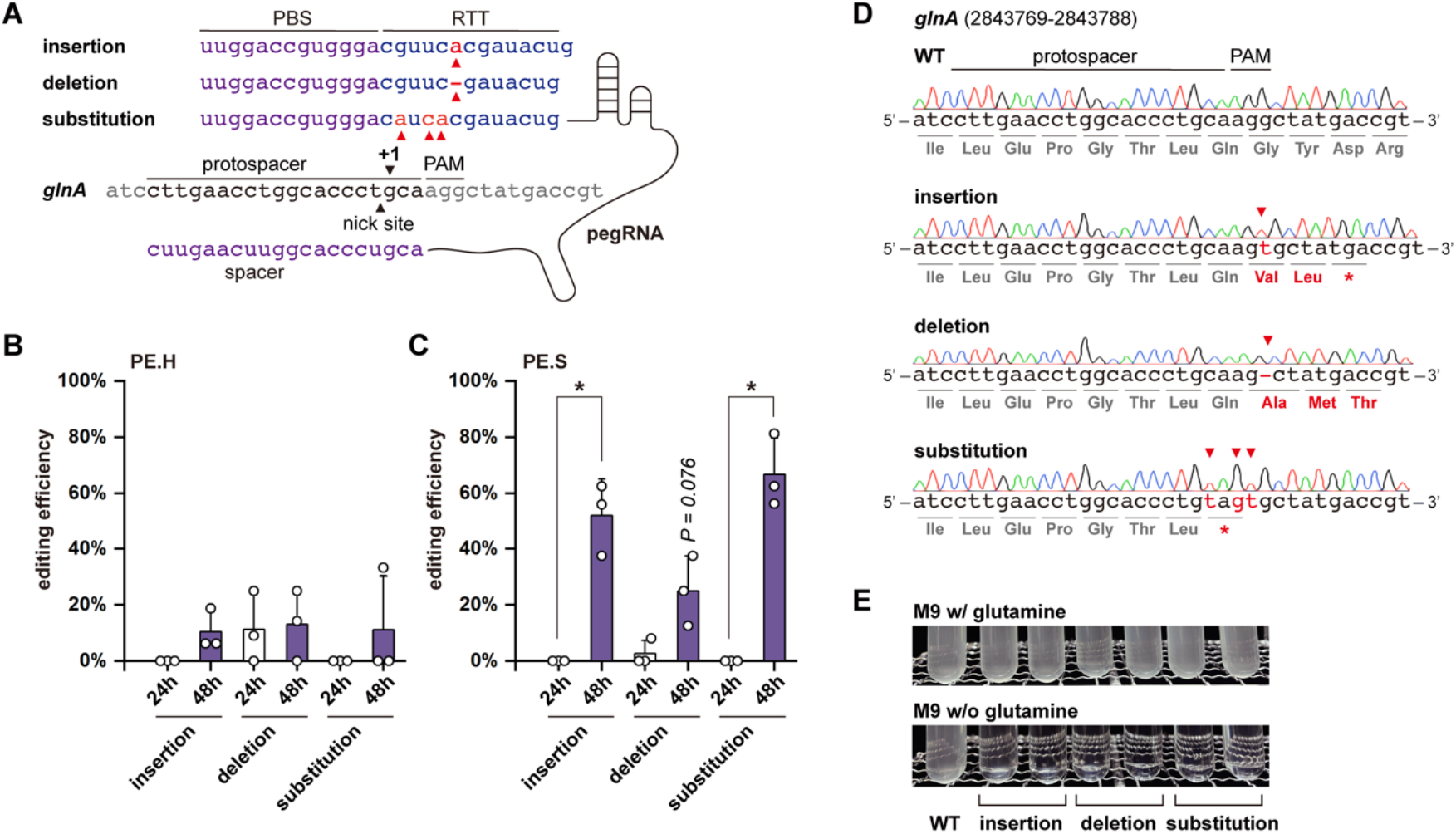
Design and demonstration of prime editing in EcN. **(A)** Design of pegRNAs. A pegRNA consists of a gRNA harboring the designed spacer, a PBS and RTT. Three pegRNAs were designed for DNA insertion (pegRNA01), deletion (pegRNA02), and substitution (pegRNA03). Prime editing efficiencies with **(B)** PE.H and **(C)** PE.S as effectors and pegRNA01, pegRNA02 and pegRNA03 for different types of editing. The editing efficiencies for 24 h induction are indicated in white columns, and those for 48 h induction were shown in purple columns. Three independent experiments were performed and the standard deviations were represented by error bars. The *t-test* was used for statistical differences evaluation (*, p < 0.05). **(D)** Sequencing results of the wild-type and edited EcN. The *glnA* was inactivated by shifting the ORF via DNA insertions and deletions or introducing premature STOP codons via substituting CAAG to TAGT. The edited positions were indicated by the red arrows, and the altered amino acids and the edited bases were highlighted in red. **(E)** Phenotypical evaluation of the *glnA* inactivated EcN.

Next, we assessed the editing efficiency at another locus in the genome and successfully achieved insertion, deletion and substitution **(Figure S2)**, illustrating the capability of our prime editing system. Notably, we found mixed sequencing signals of the edited colonies, which was due to the working mechanism of prime editing and has been reported in deamination-mediated base editing as well (14, 30). The pure edited strains can be obtained via a one-step segregation **(Figure S3)**. Finally, the working plasmids could be easily cured after one transfer in non-selective media at 37 °C **(Figure S4)**.

To verify that the expression levels of the working plasmids were indeed increased with extended induction time, we generated placI-mCherry01 **(Figure S5A and Table S1)** by replacing the editing effector with mCherry to visualize the expression levels. We observed significantly higher fluorescence of the placI-mCherry01 containing strain at 48 h than that at 24 h **(Figure S5B)**, demonstrating our assumption. Inspired by the observation, we attempted to redesign module 3 with pMB origin of replication for higher copy numbers and *sacB*, coding for the levansucrase, for plasmid curing (30) **(Figure S5C)**. However, with such a design, no detectable editing has been identified. These findings suggest that elevated expression level of the prime editing system is essential for successful prime editing, while it is not the only determining factor.

### Prime editing generates significantly fewer off-target events

Prime editing and base editing can generate off-target events in bacteria, as the system, when functioning at undesired loci, no longer kills the cell. Theoretically, prime editing generates fewer off-target events due to its multiple hybridization working mechanisms, including complementary bindings between the spacer and protospacer, the PBS and the 3’ nicked strand, as well as its specific dependence on the PAM **(Figure 3A)**. To evaluate the off-target events generated by prime editing, we whole-genome sequenced four randomly picked colonies from each edited strain with DNA insertion (PR01), deletion (PR02) and substitution (PR03) **(Figure 3A and Table S4)**. As expected, prime editing created around only 15.0 ± 3.3 off-target events, ranging from 6 to 21 **(Dataset)**. While the minimal and maximal off-target events were both identified in colonies with DNA insertions, no significant differences were observed among different editing types. Most off-target events (67.0 ± 9.0%) were located in non-coding regions or resulted in silent mutations **(Figure 3B and 3C)**. Only 4.9 ± 1.4 off-target events, ranging from 2 to 8, led to missense mutations **(Figure 3D)**, and no nonsense mutations leading to early STOP codons were identified in any of the tested colonies **(Figure 3E)**.

**Figure 3.**
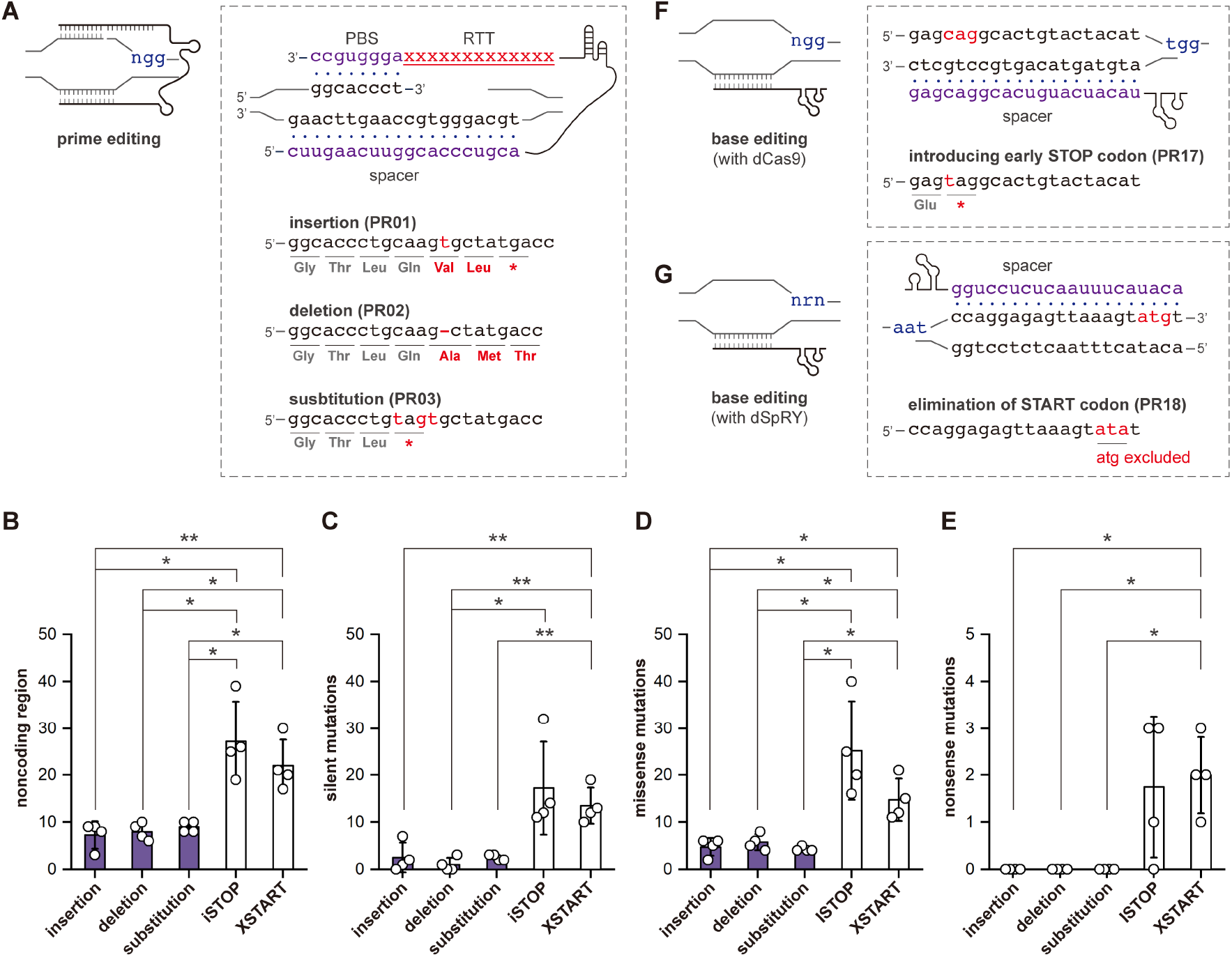
Off-target event evaluation. **(A)** Schematic illustration of the multiple hybridization mechanism of prime editing and the sequenced strains with different types of editing. The spacer and PBS, which hybridize with the target sequences are highlighted in purple. Numbers of mutations that were located in **(B)** noncoding regions, **(C)** silent mutations, **(D)** missense mutations, and **(E)** nonsense mutations. Numbers of mutations generated by prime editing and base editing are indicated in purple and white, respectively. Four colonies from each edited strain were randomly picked for whole-genome sequencing, and the standard deviations are represented by error bars. The *t-test* was used to determine the statistical differences (*, p < 0.05 and **, p < 0.01). Scheme of **(F)** iSTOP and **(G)** XSTARGT with corresponding strains PR17 and PR18, respectively.

In addition, we compared the off-target events of prime editing with those of base editing. We generated two base editors to introduce early STOP codons (iSTOP) (31, 32) and to exclude START codons (XSTART) (16) for gene inactivation **(Figure 3F and 3G)**. Using iSTOP and XSTART, we inactivated the same gene in EcN **(Figure 3F, 3G, and S6)**, and four colonies from each editing approach were sequenced. Compared to prime editing, base-edited strains showed significantly more off-target events, with iSTOP generating 71.5 ± 25.0 and XSTART generating 52.3 ± 9.8 such unintended mutations **(Dataset)**. While most occurred in non-coding regions or caused silent mutations **(Figure 3B and 3C)**, 35.0 ± 3.0% (iSTOP) and 28.0 ± 2.0% (XSTART) of these off-target events led to missense mutations **(Figure 3D)**. In addition, 7 and 8 unexpected early STOP codons were identified for iSTOP and XSTART, respectively, accounting for 2.0 ± 2.0% and 4.0 ± 2.0% of the total off-target events **(Figure 3E and Dataset)**. Despite the high efficiency of base editing, prime editing demonstrated a significant reduction in off-target events, highlighting a trade-off between efficiency and precision.

### Barcoding EcN with prime editing for strain tracking and quantification

As living diagnostics and therapeutics, it is necessary to track engineered EcN for precision medicine. We propose a novel barcoding system for EcN by inserting a short DNA sequence that, alone or in combination with the native sequence, is unique within its working microbial community. The specific DNA sequence can be introduced using our prime editing system **(Figure 4A)**. As a proof of principle, we inserted a 7-bp of DNA sequence into the coding region of *argH* **(Figure 4B)**, which encodes for the argininosuccinate lyase, generating the barcoded auxotrophic strain PR10 **(Figure S7 and Table S4)**. The combination of 7-bp insertion and an original 3-bp DNA sequence created a unique 10-bp barcode that can be exclusively identified via BLAST **(Figure 4B)**. Moreover, we established a streamlined approach to semi-quantify the EcN via Sanger sequencing of the edited region **(Figure 4C)**. First, the edited region was directly amplified from samples where tracking of engineered EcN was required. The amplicon was then Sanger sequenced, and the results, potentially a mixed sequencing signal, were analyzed using bioinformatic tools **(Figure 4C)**. To demonstrate the utility of the approach, we mixed PR10 with the wild-type *E. coli* MG1655, which harbors a very similar genomic sequence, in different ratios. We amplified the *argH* gene directly from the mixed culture, performed Sanger sequencing, and analyzed the results using TIDER **(Figure 4C)**, a bioinformatic tool that quantifies sequencing results (33). We found that the predicted abundance of EcN correlated well with the experimental values, giving a promising coefficient of determination (R^2^) of 0.989 **(Figure 4D and Table S5)**. These results suggest that the prime-editing-based barcoding system functions effectively for EcN, and the quantification can serve as a preliminary reference for EcN administration.

**Figure 4.**
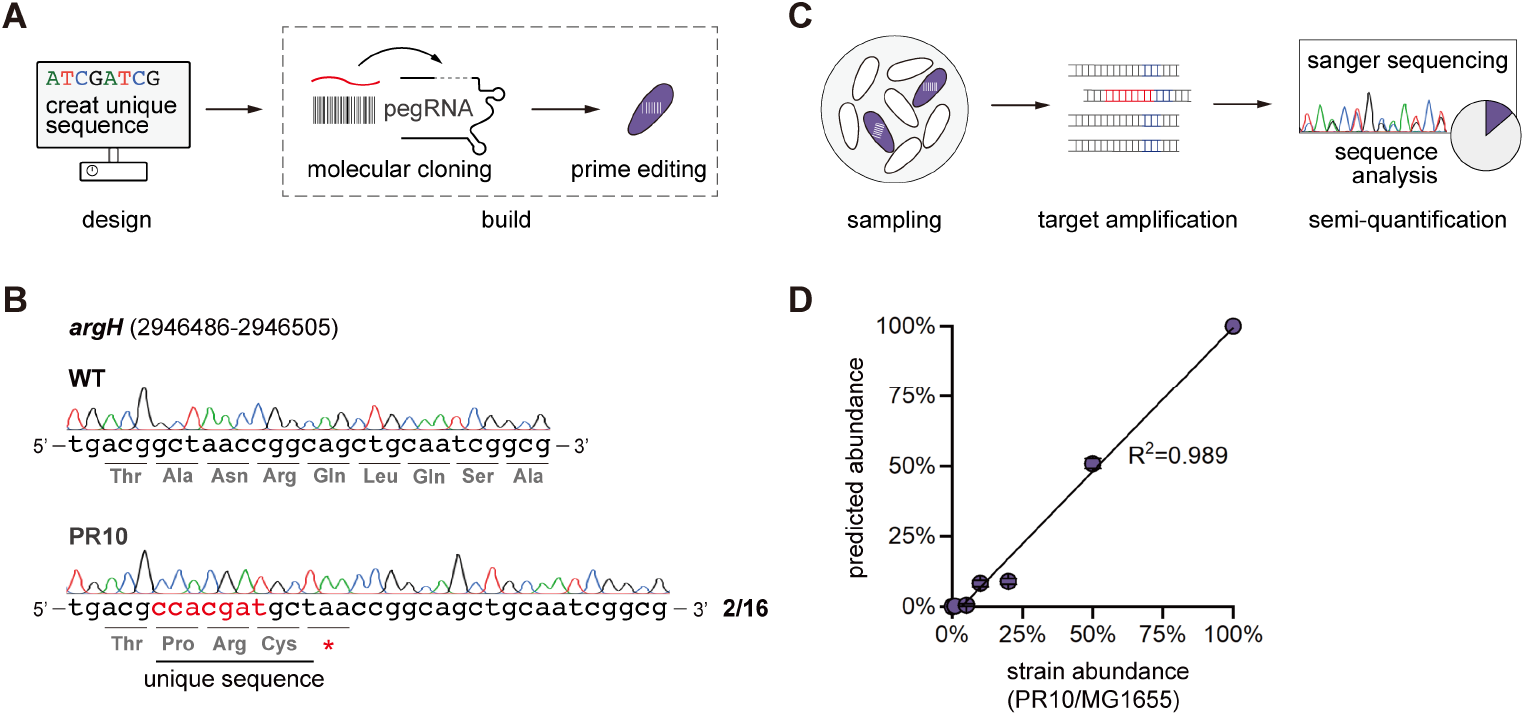
Barcoding system for strain tracking and quantification. **(A)** Schematic workflows for barcoding. Briefly, a short sequence that is unique in the working microbial community of the engineered EcN is designed in silico. Then, a customized pRC plasmid is constructed to install the barcoding sequence. **(B)** Sequencing results of the barcoded region of engineered EcN PR10. The barcoding sequence is underlined and the edited bases and the altered amino acids are highlighted in red. **(C)** Strategy for the quantification of barcoded EcN. First, the sample is taken, and PCR is then performed directly form the culture or colony. The amplicon of the edited region is Sanger sequenced and analyzed for quantification using bioinformatic tools. **(D)** Proof of principle demonstration of EcN quantification. The barcoded EcN was mixed with *E. coli* MG1655 at different but defined ratios, and the sample was directly analyzed via PCR of the liquid culture. Then the sequencing results were analyzed using TIDER. The coefficient of determination R^2^ was calculated via linear regression.

### An ARG-free platform enables non-natural functionalities

To enable non-native functionalities, engineered probiotics typically require plasmid systems, which often rely on ARGs as selection markers. However, ARGs have becoming a threatening issue to human health, and they can disseminate through horizontal gene transfer (34). Therefore, ARG-free systems are necessary for the safe engineering of probiotics. One promising strategy is the utilization of auxotrophic strains and complementary plasmids that rescue the strains from their auxotrophic deficiencies (25, 27, 35). This strategy needs a pre-engineering of auxotrophic deficiencies in the working strains. As such, we designed and built such an ARG-free platform for EcN with prime editing. First, we employed two prime-edited strains PR01 and PR10, which were deficient in glutamine and arginine **(Figure S8 and Table S4)**, respectively, as the auxotrophic hosts. Two complementary plasmids were constructed to enable non-native functionalities. The first plasmid, pRED, contains the complementary *glnA*, a pBBR1 origin of replication, and *mCherry* as the GOI **(Figure 5A and Figure S9)**. The second plasmid, pGREEN, consists of *argH*, a pMB origin of replication, and *gfp* as the GOI **(Figure 5A and Figure S9)**. Notably, we deployed codon-modified versions of *glnA* (*glnA*^*m*^) and *argH* (*argH*^*m*^) to avoid the recovery of the wild-type genes in the genome through homology-directed recombination **(Figure 5A and Table S2)**. Moreover, we used prime editing to generate a double-auxotrophic EcN strain, PR13 **(Figure S10 and Table S4)**, with both glutamine and arginine deficiencies **(Figure S11)**. PR13 was able to harbor pRED and pGREEN at the same time, enabling possibilities for more sophisticated biological tasks.

**Figure 5.**
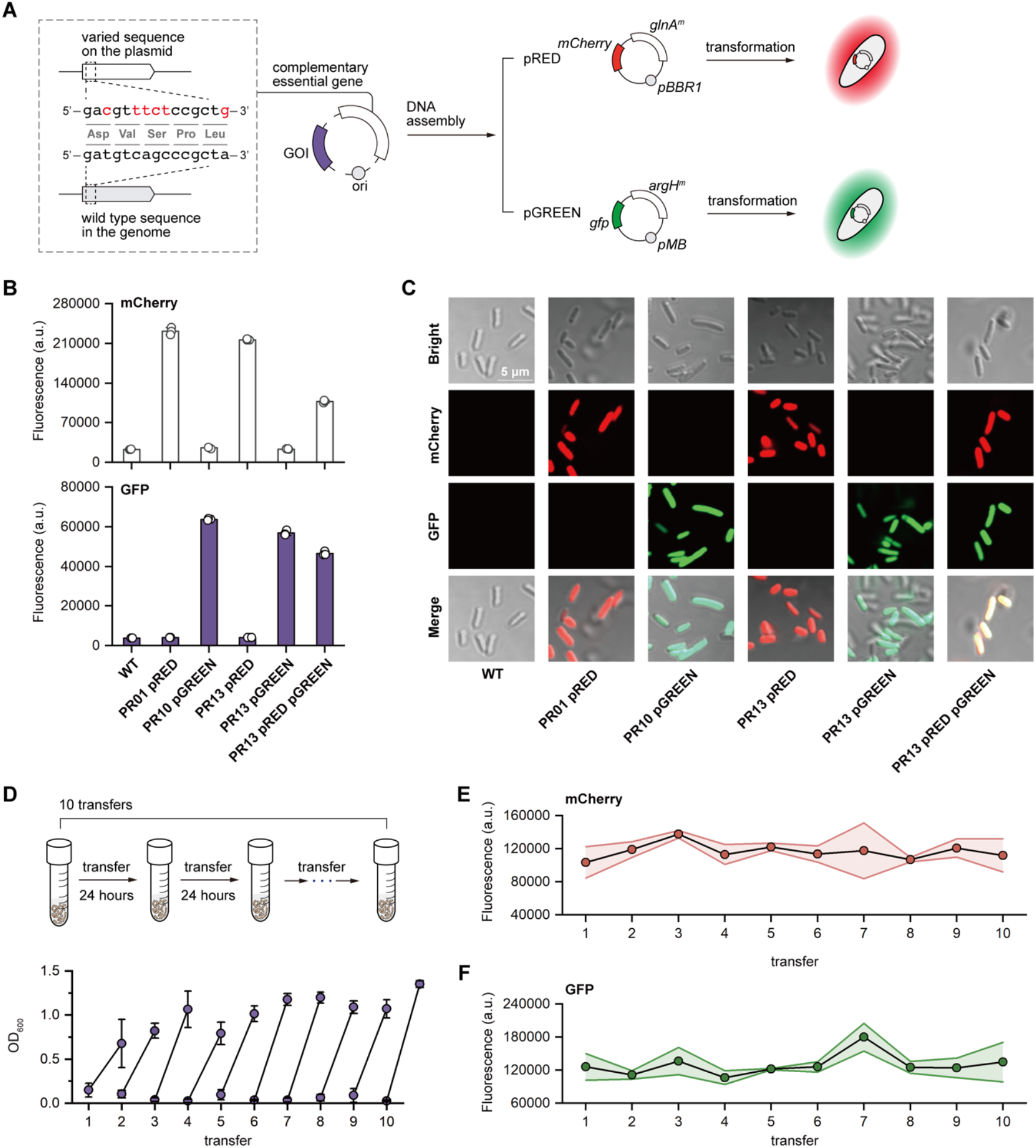
Design and evaluation of the ARG-free platform. **(A)** The ARG-free plasmids consist of the origin of replication, the complementary essential genes and GOIs. We deploy codon-modified versions of essential genes without changing the amino acids, as represented, to minimize the possibility of homology-directed recombination and the recovery of the wild-type essential genes. As proof of principle, we generated pRED with *glnA*^*m*^ (codon-modified *glnA*) and *mCherry* as the GOI, and constructed pGREEN with *argH*^*m*^ (codin-modified *argH*) and *gfp* as GOI. Then, pRED and pGREEN were transformed into PR01 and PR10, respectively for evaluation. **(B)** Fluorescence measurement of the ARG-free platform. **(C)** Visualization of the ARG-free platform in auxotrophic EcN strains by CLSM imaging. **(D)** Serial transfer experiment of the double-auxotrophic PR13 with both pRED and pGREEN. The growth profiles were measured by OD_600_. The transfer experiments were performed in triplicate, and the error bars represent the standard deviations. Fluorescence intensity of **(E)** mCherry and **(F)** GFP in double-auxotrophic EcN during the transfer experiments.

As designed, we observed that PR01 with pRED could grow in M9 minimal medium without glutamine, and PR10 with pGREEN could grow without arginine. To the contrary, PR01 and PR10 alone could not grow in the corresponding amino acid depleted media **(Figure S8)**. Furthermore, PR13 with both pRED and pGREEN was able to grow without glutamine and arginine, while PR13 carrying one of the plasmids could survive in the corresponding amino acid depleted medium **(Figure S11)**, demonstrating the effectiveness of our ARG-free plasmids for selection and maintenance. Next, we measured and visualized the fluorescence of the strains with pRED and pGREEN **(Figure 5B and 5C)**. All strains with plasmids exhibited designed fluorescence signals, which were clearly identified through confocal laser scanning microscopy (CLSM) **(Figure 5C)**. Notably, PR13 with pRED and pGREEN showed both red and green fluorescence, although the intensity of each was slightly weaker than those expressed alone **(Figure 5B)**. Finally, we evaluated the stability of our system in the double-auxotrophic PR13. We performed a serial transfer experiment of PR13 with both pRED and pGREEN, transferring the strain every 24 h for a total of 240 h, corresponding to 42 generations **(Figure 5D)**. The growth profiles were similar across transfers **(Figure 5D)**, and, despite some fluctuations, the red and green fluorescence remained stable throughout the experiment **(Figure 5E and 5F)**. These results demonstrated that our prime edited strains and the ARG-free plasmids have the potential to expand the spectrum of functionalities that engineered probiotics can perform.

## Discussion

Prime editing couples the CRISPR-Cas system with reverse transcription for precision targeting and versatile DNA modifications. One common barrier that limits its application is the relatively low editing efficiency (19, 21). Efforts have been made to enhance prime editing efficiency, such as mining or engineering new effectors (e.g., RT) (36-38), combining extra genetic modules (39-41), and designing customized pegRNAs (42-44). However, it appears more challenging in bacteria, where only a few reports have demonstrated the utilization of prime editing in bacterial hosts, including *E. coli* (20, 21), pathogenic *Streptococcus pneumoniae* (22), *Leptospira borgpetersenii* (23) and *Klebsiella pneumoniae* (21), with efficiency still requiring improvement or necessitating pre-engineering of the target strain. Here, we present a straightforward prime editing approach with only essential genetic modules and an optimized workflow, showing promising editing performance. This was achieved by simply selecting a suitable editing effector, the CRISPR-Cas system from *S. pyogenes* with its native codons and codon-optimized RT, as well as extending the induction time to 48 h. All types of prime editing were successfully achieved in every individual round of experiments with promising efficiencies **(Figure 2B and 2C)**. We acknowledge that the efficiency of prime editing fluctuates and needs further improvement, while this may represent an unavoidable trade-off between efficiency, versatility and precision.

Expanding prime editing to bacteria does not seem urgently required, especially for model strains (e.g., *E. coli* MG1655), where a plethora of gene editing tools have already been developed or adapted. Even for non-model strains, where conventional CRISPR-Cas systems encounter challenges, base editing has been established for gene inactivation in various bacterial species with an overall efficiency reaching 100% (7, 16). However, we noticed limitations in base editing, including the constrained editing spectrum and the inevitable off-target effects that traded for high efficiency (16, 17, 45). While the edited strains may be acceptable for industrial applications after careful evaluation, they may raise concerns in medical contexts. Precision engineering of probiotic bacteria requires minimal undesired interferences in the genome to promote human health and treat diseases. At this moment, prime editing finds its unique niche. Using prime editing, we successfully established a barcoding system for strain tracking and an ARG-free platform to enable non-natural functionalities, underscoring its versatile potential in clinical applications. Meanwhile, we observed only a few unintended mutations in the edited strains, which is more obvious when compared with two distinct base editing strategies **(Figure 3)**, emphasizing its superior precision in genome editing. These advantages make prime editing an ideal methodology to fully unveil the capability of engineered probiotics via either intensive strain engineering or the interrogation of cellular physiology.

CRISPR-Cas systems are renowned for precision and versatility as genome editing tools, owing to a streamlined working machinery. This is especially critical for probiotic bacteria, which are often nature-isolated strains that present unique challenges in genetic manipulation. As such, we deliberately minimized the complexity of our system, awakening a back-to-basics CRISPR-Cas gene editing strategy without sophisticated and extraneous designs, which can be readily implemented in any laboratory with the basic CRISPR setups. We believe our work provides not only a powerful gene editing approach for precision medicine but also a paradigm for developing advanced, yet accessible, CRISPR-Cas systems.

## Methods

### Strains and media

All the strains used in this study are listed in **Table S4**, and all edited strains were plasmid-cured before evaluation. *E. coli* DH5α (Takara Bio) was used for general molecular cloning. EcN was a generous gift from Dr. Chun Loong Ho. *E. coli* were cultivated at 37 °C in LB medium which is composed of 1% NaCl, 1% tryptone, and 0.5% yeast extract (solid medium with 1.5% agar). For phenotypic analysis, the wild-type and edited EcN were grown in M9 minimal medium containing 15.14 g/L Na_2_HPO_4_×12H_2_O, 3.0 g/L KH_2_PO_4_, 0.5 g/L NaCl, 1.0 g/L NH_4_Cl, 0.241 g/L MgSO_4_, 0.011 g/L CaCl_2_ and 4 g/L glucose. Appropriate antibiotics (100 µg mL^−1^ ampicillin and 20 µg mL^−1^ gentamicin) and amino acids (5 mM L-glutamine and 0.4 g/L L-arginine) were supplemented when required. The IPTG was added for induction with the final concentration of 1 mM.

### Plasmid construction

All of the DNA manipulation and molecular cloning were based on the following protocols unless otherwise noted. DNA fragments were amplified by PCR using PrimeSTAR^®^ Max DNA Polymerase (Takara Bio), and plasmids were generated via DNA assembly strategy using In-Fusion^®^ Snap Assembly Master Mix (Takara Bio). Plasmids were extracted by QIAprep Spin Miniprep Kit (Qiagen), quantified by NanoDrop One (Thermo Scientific) and confirmed by gel electrophoresis and Sanger sequencing. All the plasmids used in this study are summarized in **Table S1**, and the sequences of the codon-optimized RT and the codon-modified essential genes (*glnA*^*m*^ and *argH*^*m*^) are listed in **Table S2**. All pegRNAs in this work are given in **Table S3**. The primers used in this work are listed in **Table S6**.

To construct the working plasmids of prime editing, we assembled the PE2 from pCMV-PE2-P2A-GFP, the *E. coli* codon-optimized RT, and *oriR101* from pKD46 to the plasmid pWY, replacing the base editor and generating pPE.H. Then, we replaced PE.H with PE.S, *ncas9* (H840A) from *S. pyogenes* fused with RT, generating pPE.S. The pegRNA cassettes were synthesized and implemented in pPE.H or pPE.S to generate pRC serial working plasmids. To generate placl-mCherry01, *mCherry* from pmCherry was assembled into pRC02 to substitute PE.H. The levansucrase coding gene, *sacB*, and *pMB* ori were cloned into placI-mCherry01, exchanging *oriR101* and generating placI-mCherry02. pRED was built by integrating *glnA*^*m*^ to replace the *GmR* in pmCherry. pGREEN was built by assembling *gfp* from pAM4787 and *argH*^*m*^ with pBR322 as the backbone. To construct the base editing plasmid, gRNA-stop for installing early STOP codon in and gRNA-start for eliminating START codon of *glnA* were integrated in pBeCas9 and pBeSpRY (16), generating pBeCas9-iSTOP and pBeSpRY-XSTART.

### Transformation and prime editing of *E. coli* Nissle 1917

To prepare the chemically competent cells, *E. coli* Nissle 1917 was cultivated in LB liquid medium at 37 °C and 180 rpm. When OD_600_ reached 0.3 - 0.5, the cells were harvested at 4 °C and 5000 rpm for 10 min. The pellets were washed twice using 0.1 M CaCl_2_, and resuspended and aliquoted in 0.1 M CaCl_2_ and 15% (v/v) of glycerol for storage. For transformation, the working plasmids (25 ng) were added to competent cells and kept on ice for 20 min, followed by heat shock at 42 °C for 60 s. Cells were recovered in 1 mL of LB medium at 30 °C for 1 h and then plated on LB agar plates with appropriate antibiotics or undergone subsequent prime editing procedures. For prime editing, we used the optimized liquid-induction approach established in our previous work (7). After transformation, IPTG (1 mM) and gentamicin (20 µg mL^−1^) were added to the recovered mixture for induction. Then, the cells were plated on LB agar plates with gentamicin to obtain transformants. The transformants were screened for successfully edited strains via Sanger sequencing the editing region in the genome.

### Plasmid curing

The edited strain was cultivated in LB liquid medium at 37 °C without antibiotics for 14 to 16 h, and streaked on LB agar plates to obtain single colonies. The curing of plasmids was determined by the failed amplification of specific DNA fragments in the plasmids and recovery of sensitivity to gentamicin.

### Whole-genome sequencing and analysis

To evaluate the off-target events of prime editing and base editing, the wild-type and engineered EcN were whole-genome sequenced and analyzed. For the edited strains, four independent colonies were randomly selected, and the whole genome sequencing and off-target effect analysis were performed following established procedures in our laboratory (7, 15). The genome sequence with NCBI accession number NZ_CP082949.1 was employed as a reference, while the results of edited strains were compared with that of the wild-type EcN. The sequencing data are stored in NCBI with the accession number PRJNA1136227.

### Quantification of barcoded EcN

The mixtures of wild-type *E. coli* MG1655 and PR10 were harvested and washed twice before analysis. Then, we amplified the barcoded regions of each mixed culture for Sanger sequencing. The sequencing results were analyzed by TIDER following the suggested instructions (https://tide.nki.nl) (33).

### Fluorescence measurement and visualization

To obtain the fluorescence intensities, we washed the cells twice with phosphate buffered saline, and, after measured the optical density at 600 nm, we diluted the cells to OD_600_ of 0.01 for the measurement using a microplate reader (TECAN). The excitation and the emission wavelength for mCherry were 590 nm and 645 nm, respectively, and the excitation and the emission wavelength for GFP were 488 nm and 530 nm. For visualization, the cells were harvested and washed using the same protocol as above, and visualized using CLSM (Zeiss) with excitation wavelengths of 561 nm and 488 nm for mCherry and GFP, respectively.

### Serial transfer experiment

The evaluate the stability of our ARG-free platform, serial transfer experiment was performed at 37 °C and 180 rpm in M9 minimal medium without amino acid supplements. After every 24 h of cultivation, the culture was transferred into fresh M9 medium and cultivated under the same condition. Samples from the previous culture and new transfer were collected at inoculation and measured for fluorescence and OD_600_. The number of generations (n) was calculated by the following equation:

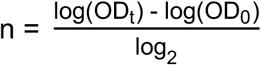

where OD_t_ is the OD_600_ of each transfer after 24 h cultivation, and OD_0_ is the initial OD_600_ of each transfer. The total number of generations is the sum of the generation number of each transfer (6).

## Supporting information

Dataset of off-target events

## Conflict of Interests

The authors declare no conflict of interest.

## Acknowledgments

This work was supported by the National Natural Science Foundation of China (22278246, U20A20146 and 22378233), the Department of Science and Technology of Shandong Province (2022HWYQ-017), the Natural Science Foundation of Shandong Province (ZR2021ME066) and the Qilu Young Scholar Program of Shandong University (to P.-F.X.), and the Taishan Scholars Project of Shandong Province (NO. tstp20230604).

## Notes

### Competing Interest Statement

The authors have declared no competing interest.

## References

1. D. Collias et al., Systematically attenuating DNA targeting enables CRISPR-driven editing in bacteria. Nat. Commun. 14, 680 (2023).

2. J. M. Vento, N. Crook, C. L. Beisel, Barriers to genome editing with CRISPR in bacteria. J. Ind. Microbiol. Biotechonl. 46, 1327–1341 (2019).

3. D. T. Riglar, P. A. Silver, Engineering bacteria for diagnostic and therapeutic applications. Nat. Rev. Microbiol. 16, 214–225 (2018).

4. M. Pan et al., Genomic and epigenetic landscapes drive CRISPR-based genome editing in Bifidobacterium. Proc. Natl. Acad. Sci. U. S. A. 119, e2205068119 (2022).

5. P. F. Xia, H. Ling, J. L. Foo, M. W. Chang, Synthetic biology toolkits for metabolic engineering of cyanobacteria. Biotechnol. J. 14, e1800496 (2019).

6. P. F. Xia et al., Reprogramming acetogenic bacteria with CRISPR-targeted base editing via deamination. ACS Synth. Biol. 9, 2162–2171 (2020).

7. Y. Wei, L. J. Feng, X. Z. Yuan, S. G. Wang, P. F. Xia, Developing a base editing system for marine Roseobacter clade bacteria. ACS Synth. Biol. 12, 2178–2186 (2023).

8. Y. Wei, S. G. Wang, P. F. Xia, Blue valorization of lignin-derived monomers via reprogramming marine bacterium Roseovarius nubinhibens. Appl. Environ. Microbiol. 90, e0089024 (2024).

9. A. V. Anzalone, L. W. Koblan, D. R. Liu, Genome editing with CRISPR-Cas nucleases, base editors, transposases and prime editors. Nat. Biotechnol. 38, 824–844 (2020).

10. M. Zhang, Z. Zhu, G. Xun, H. Zhao, To cut or not to cut: next-generation genome editors for precision genome engineering. Curr. Opin. Biomed. Eng. 28, 100489 (2023).

11. A. V. Anzalone et al., Search-and-replace genome editing without double-strand breaks or donor DNA. Nature 576, 149–157 (2019).

12. A. C. Komor, Y. B. Kim, M. S. Packer, J. A. Zuris, D. R. Liu, Programmable editing of a target base in genomic DNA without double-stranded DNA cleavage. Nature 533, 420–424 (2016).

13. K. Nishida et al., Targeted nucleotide editing using hybrid prokaryotic and vertebrate adaptive immune systems. Science 353 (2016).

14. Y. Tong et al., Highly efficient DSB-free base editing for streptomycetes with CRISPR-BEST. Proc. Natl. Acad. Sci. U. S. A. 116, 20366–20375 (2019).

15. S. Y. Wang, X. Li, S. G. Wang, P. F. Xia, Base editing for reprogramming cyanobacterium Synechococcus elongatus. Metab. Eng. 75, 91–99 (2023).

16. X. Li, Y. Wei, S. Y. Wang, S. G. Wang, P. F. Xia, One-for-all gene inactivation via PAM-independent base editing in bacteria. bioRxiv [preprint] (2024). 10.1101/2024.06.17.599441 (accessed 15 October 2024).

17. F. M. Seys et al., Base editing enables duplex point mutagenesis in Clostridium autoethanogenum at the price of numerous off-target mutations. Front. Bioeng. Biotechnol. 11, 1211197 (2023).

18. C. M. Whitford et al., Systems analysis of highly multiplexed CRISPR-base editing in streptomycetes. ACS Synth. Biol. 12, 2353–2366 (2023).

19. P. J. Chen, D. R. Liu, Prime editing for precise and highly versatile genome manipulation. Nat. Rev. Genet. 24, 161–177 (2023).

20. Y. Tong, T. S. Jorgensen, C. M. Whitford, T. Weber, S. Y. Lee, A versatile genetic engineering toolkit for E. coli based on CRISPR-prime editing. Nat. Commun. 12, 5206 (2021).

21. H. Zhang et al., BacPE: a versatile prime-editing platform in bacteria by inhibiting DNA exonucleases. Nat. Commun. 15, 825 (2024).

22. M. Rengifo-Gonzalez et al., Make-or-break prime editing for bacterial genome engineering. bioRxiv [preprint] (2024). 10.1101/2024.06.27.601116 (accessed 15 October 2024).

23. L. G. V. Fernandes et al., CRISPR-prime editing, a versatile genetic tool to create specific mutations with a single nucleotide resolution in Leptospira. mBio 15, e0151624 (2024).

24. K. J. Chua, W. C. Kwok, N. Aggarwal, T. Sun, M. W. Chang, Designer probiotics for the prevention and treatment of human diseases. Curr. Opin. Chem. Biol. 40, 8–16 (2017).

25. I. Y. Hwang et al., Engineered probiotic Escherichia coli can eliminate and prevent Pseudomonas aeruginosa gut infection in animal models. Nat. Commun. 8, 15028 (2017).

26. E. Koh et al., Engineering probiotics to inhibit Clostridioides difficile infection by dynamic regulation of intestinal metabolism. Nat. Commun. 13, 3834 (2022).

27. C. L. Ho et al., Engineered commensal microbes for diet-mediated colorectal-cancer chemoprevention. Nat. Biomed. Eng. 2, 27–37 (2018).

28. C. R. Gurbatri et al., Engineering tumor-colonizing E. coli Nissle 1917 for detection and treatment of colorectal neoplasia. Nat. Commun. 15, 646 (2024).

29. A. J. Triassi et al., Redesign of an Escherichia coli Nissle treatment for phenylketonuria using insulated genomic landing pads and genetic circuits to reduce burden. Cell Syst. 14, 512–524 e512 (2023).

30. S. D. Rodrigues et al., Efficient CRISPR-mediated base editing in Agrobacterium spp. Proc. Natl. Acad. Sci. U. S. A. 118, e2013338118 (2021).

31. P. Billon et al., CRISPR-mediated base editing enables efficient disruption of eukaryotic genes through induction of STOP codons. Mol. Cell 67, 1068–1079 e1064 (2017).

32. C. Kuscu et al., CRISPR-STOP: gene silencing through base-editing-induced nonsense mutations. Nat. Methods 14, 710–712 (2017).

33. E. K. Brinkman et al., Easy quantification of template-directed CRISPR/Cas9 editing. Nucleic Acids Res. 46, e58 (2018).

34. D. G. J. Larsson, C. F. Flach, Antibiotic resistance in the environment. Nat. Rev. Microbiol. 20, 257–269 (2022).

35. M. B. Amrofell et al., Engineering E. coli strains using antibiotic-resistance-gene-free plasmids. Cell Rep. Methods 3, 100669 (2023).

36. Z. Gao et al., A truncated reverse transcriptase enhances prime editing by split AAV vectors. Mol. Ther. 30, 2942–2951 (2022).

37. C. Zheng et al., A flexible split prime editor using truncated reverse transcriptase improves dual-AAV delivery in mouse liver. Mol. Ther. 30, 1343–1351 (2022).

38. J. Grunewald et al., Engineered CRISPR prime editors with compact, untethered reverse transcriptases. Nat. Biotechnol. 41, 337–343 (2023).

39. P. J. Chen et al., Enhanced prime editing systems by manipulating cellular determinants of editing outcomes. Cell 184, 5635–5652 e5629 (2021).

40. J. Ferreira da Silva et al., Click editing enables programmable genome writing using DNA polymerases and HUH endonucleases. Nat. Biotechnol. 10.1038/s41587-024-02324-x (2024).

41. P. Liu et al., Increasing intracellular dNTP levels improves prime editing efficiency. Nat. Biotechnol. 10.1038/s41587-024-02405-x (2024).

42. J. W. Nelson et al., Engineered pegRNAs improve prime editing efficiency. Nat. Biotechnol. 40, 402–410 (2022).

43. X. Li et al., Highly efficient prime editing by introducing same-sense mutations in pegRNA or stabilizing its structure. Nat. Commun. 13, 1669 (2022).

44. X. Lei et al., Rapid generation of long, chemically modified pegRNAs for prime editing. Nat. Biotechnol. 10.1038/s41587-024-02394-x (2024).

45. E. Kozaeva, Z. S. Nielsen, M. Nieto-Dominguez, P. I. Nikel, The pAblo.pCasso self-curing vector toolset for unconstrained cytidine and adenine base-editing in Gramnegative bacteria. Nucleic Acids Res 52, e19 (2024).

