## Supporting information for "Awakening prime editing for precision engineering of probiotic *Escherichia coli* Nissle 1917"

* Correspondence:

Peng-Fei Xia

**Table S1.** Plasmids used in this study.

| Name | Description | Source |
|---|---|---|
| pBR322 | *colE1 ori, bla, tetA* | Lab stock |
| pRE118 | *R6K γ* ori, *sacB*, *kanR* | Lab stock |
| pKD46 | *oriR101*, *bla* | Lab stock (1) |
| pBBR1MCS-5 | *pBBR1* ori, *GmR* | Lab stock (2) |
| pAM4787 | *ColE1 ori*, partial sequence from pANS | Addgene #120088 (3) |
| pCMV-PE2-P2A-GFP | Cas9 (H840A)-M-MLVrt (D200N, T306K, W313F, T330P, L603W) | Addgene #132776 (4) |
| pWY | pBBR1MCS-5, *lacI*-$P_{trc}$, *dcas9*, *PmCDA1*, *ugi* | Lab stock (2) |
| pmCherry | pBBR1MCS-5, *lacI*-$P_{trc}$, *mCherry* | Lab stock (2) |
| pBeSpRY | *oriR101*, *bla*, *trc* promoter, *dSpRY-PmCDA1-ugi* | Lab stock (5) |
| pBeCas9 | *oriR101*, *bla*, *trc* promoter, *dCas9-PmCDA1-ugi* | Lab stock (5) |
| pPegRNA01 | pUC57, pegRNA01 | This study |
| pPegRNA02 | pUC57, pegRNA02 | This study |
| pPegRNA03 | pUC57, pegRNA03 | This study |
| pPegRNA04 | pUC57, pegRNA04 | This study |
| pPegRNA05 | pUC57, pegRNA05 | This study |
| pPegRNA06 | pUC57, pegRNA06 | This study |
| pPegRNA07 | pUC57, pegRNA07 | This study |
| pgRNA-stop | pUC57, gRNA01 | This study |
| pgRNA-start | pUC57, gRNA02 | This study |
| pPE.H | PE.H (*ncas9* from pCMV-PE2-P2A-GFP fused with *E. coli* codon-optimized RT), *lacI*-$P_{trc}$, *GmR*, *oriR101* | This study |
| pPE.S | PE.S (*ncas9* from *S. pyogenes* with native sequence fused with *E. coli* codon-optimized RT), *lacI*-$P_{trc}$, *GmR*, *oriR101* | This study |

| | | |
|---|---|---|
| pRC01 | pPE.H, pegRNA01 | This study |
| pRC02 | pPE.H, pegRNA02 | This study |
| pRC03 | pPE.H, pegRNA03 | This study |
| pRC04 | pPE.S, pegRNA01 | This study |
| pRC05 | pPE.S, pegRNA02 | This study |
| pRC06 | pPE.S, pegRNA03 | This study |
| pRC07 | pPE.H, pegRNA04 | This study |
| pRC08 | pPE.H, pegRNA05 | This study |
| pRC09 | pPE.H, pegRNA06 | This study |
| pRC10 | pPE.H, pegRNA07 | This study |
| placI-mCherry01 | pRC02, PE.H::*mcherry, lacI*-$P_{trc}$, *GmR*, *oriR101* | This study |
| placI-mCherry02 | placI-mCherry01, *oriR101*::*pMB* ori*, sacB* | This study |
| pRED | pmCherry, *GmR*::$glnA^{m}$ | This study |
| pGREEN | pBR322, *gfp*, $argH^{m}$ | This study |
| pBeCas9-iSTOP | pBeCas9, gRNA-stop | This study |
| pBeSpRY-XSTART | pBeSpRY, gRNA-start | This study |

**Table S2.** Sequences of codon-optimized RT and *glnA*$^{m}$ and *argH*$^{m}$.

| **Gene** | **Sequence** |
|---|---|
| codon-optimized RT | ACCCTGAACATCGAAGACGAATACCGTCTGCACGAAACCTCTAAA<br>GAACCGGACGTTTCTCTGGGTTCTACCTGGCTGTCTGACTTCCCG<br>CAGGCTTGGGCTGAAACCGGTGGTATGGGTCTGGCTGTTCGTCA<br>GGCTCCGCTGATCATCCCGCTGAAAGCTACCTCTACCCCGGTTTC<br>TATCAAACAGTACCCGATGTCTCAGGAAGCTCGTCTGGGTATCAA<br>ACCGCACATCCAGCGTCTGCTGGACCAGGGTATCCTGGTTCCGT<br>GCCAGTCTCCGTGGAACACCCCGCTGCTGCCGGTTAAAAAACCG<br>GGTACCAACGACTACCGTCCGGTTCAGGACCTGCGTGAAGTTAAC<br>AAACGTGTTGAAGACATCCACCCGACCGTTCCGAACCCGTACAAC<br>CTGCTGTCTGGTCTGCCGCCGTCTCACCAGTGGTACACCGTTCTG<br>GACCTGAAAGACGCTTTCTTCTGCCTGCGTCTGCACCCGACCTCT<br>CAGCCGCTGTTCGCTTTCGAATGGCGTGACCCGGAAATGGGTATC<br>TCTGGTCAGCTGACCTGGACCCGTCTGCCGCAGGGTTTCAAAAAC<br>TCTCCGACCCTGTTCAACGAAGCTCTGCACCGTGACCTGGCTGAC<br>TTCCGTATCCAGCACCCGGACCTGATCCTGCTGCAGTACGTTGAC<br>GACCTGCTGCTGGCTGCTACCTCTGAACTGGACTGCCAGCAGGG<br>TACCCGTGCTCTGCTGCAGACCCTGGGTAACCTGGGTTACCGTGC<br>TTCTGCTAAAAAAGCTCAGATCTGCCAGAAACAGGTTAAATACCTG<br>GGTTACCTGCTGAAAGAAGGTCAGCGTTGGCTGACCGAAGCTCGT<br>AAAGAAACCGTTATGGGTCAGCCGACCCCGAAAACCCCGCGTCA<br>GCTGCGTGAATTCCTGGGTAAAGCTGGTTTCTGCCGTCTGTTCAT<br>CCCGGGTTTCGCTGAAATGGCTGCTCCGCTGTACCCGCTGACCAA<br>ACCGGGTACCCTGTTCAACTGGGGTCCGGACCAGCAGAAAGCTT<br>ACCAGGAAATCAAACAGGCTCTGCTGACCGCTCCGGCTCTGGGT<br>CTGCCGGACCTGACCAAACCGTTCGAACTGTTCGTTGACGAAAAA<br>CAGGGTTACGCTAAAGGTGTTCTGACCCAGAAACTGGGTCCGTGG<br>CGTCGTCCGGTTGCTTACCTGTCTAAAAAACTGGACCCGGTTGCT<br>GCTGGTTGGCCGCCGTGCCTGCGTATGGTTGCTGCTATCGCTGTT<br>CTGACCAAAGACGCTGGTAAACTGACCATGGGTCAGCCGCTGGTT<br>ATCCTGGCTCCGCACGCTGTTGAAGCTCTGGTTAAACAGCCGCCG<br>GACCGTTGGCTGTCTAACGCTCGTATGACCCACTACCAGGCTCTG<br>CTGCTGGACACCGACCGTGTTCAGTTCGGTCCGGTTGTTGCTCTG<br>AACCCGGCTACCCTGCTGCCGCTGCCGGAAGAAGGTCTGCAGCA<br>CAACTGCCTGGACATCCTGGCTGAAGCTCACGGTACCCGTCCGG<br>ACCTGACCGACCAGCCGCTGCCGGACGCTGACCACACCTGGTAC<br>ACCGACGGTTCTTCTCTGCTGCAGGAAGGTCAGCGTAAAGCTGGT<br>GCTGCTGTTACCACCGAAACCGAAGTTATCTGGGCTAAAGCTCTG<br>CCGGCTGGTACCTCTGCTCAGCGTGCTGAACTGATCGCTCTGACC<br>CAGGCTCTGAAAATGGCTGAAGGTAAAAAACTGAACGTTTACACC<br>GACTCTCGTTACGCTTTCGCTACCGCTCACATCCACGGTGAAATC<br>TACCGTCGTCGTGGTTGGCTGACCTCTGAAGGTAAAGAAATCAAA<br>AACAAAGACGAAATCCTGGCTCTGCTGAAAGCTCTGTTCCTGCCG<br>AAACGTCTGTCTATCATCCACTGCCCGGGTCACCAGAAAGGTCAC<br>TCTGCTGAAGCTCGTGGTAACCGTATGGCTGACCAGGCTGCTCGT<br>AAAGCTGCTATCACCGAAACCCCGGACACCTCTACCCTGCTGATC |

GAAAACTCTTCTCCGTCTGGTGGTTCTAAACGTACCGCTGACGGT
TCTGAATTCGAATGA

*glnA*$^{m}$

ATGTCTGCTGAACACGTTCTGACCATGCTGAACGAACACGAAGTT
AAATTCGTTGACCTGCGTTTCACCGACACCAAAGGTAAAGAACAG
CACGTTACCATCCCGGCTCACCAGGTTAACGCTGAATTCTTCGAA
GAAGGTAAAATGTTCGACGGTTCTTCTATCGGTGGTTGGAAAGGT
ATCAACGAATCTGACATGGTTCTGATGCCGGACGCTTCTACCGCT
GTTATCGACCCGTTCTTCGCTGACTCTACCCTGATCATCCGTTGC
GACATCCTGGAACCGGGTACCCTGCAGGGTTACGACCGTGACCC
GCGTTCTATCGCTAAACGTGCTGAAGACTACCTGCGTTCTACCGG
TATCGCTGACACCGTTCTGTTCGGTCCGGAACCGGAATTCTTCCT
GTTCGACGACATCCGTTTCGGTTCTTCTATCTCTGGTTCTCACGTT
GCTATCGACGACATCGAAGGTGCTTGGAACTCTTCTACCCAGTAC
GAAGGTGGTAACAAAGGTCACCGTCCGGCTGTTAAAGGTGGTTAC
TTCCCGGTTCCGCCGGTTGACTCTGCTCAGGACATCCGTTCTGAA
ATGTGCCTGGTTATGGAACAGATGGGTCTGGTTGTTGAAGCTCAC
CACCACGAAGTTGCTACCGCTGGTCAGAACGAAGTTGCTACCCGT
TTCAACACCATGACCAAAAAAGCTGACGAAATCCAGATCTACAAAT
ACGTTGTTCACAACGTTGCTCACCGTTTCGGTAAAACCGCTACCTT
CATGCCGAAACCGATGTTCGGTGACAACGGTTCTGGTATGCACTG
CCACATGTCTCTGTCTAAAAACGGTGTTAACCTGTTCGCTGGTGAC
AAATACGCTGGTCTGTCTGAACAGGCTCTGTACTACATCGGTGGT
GTTATCAAACACGCTAAAGCTATCAACGCTCTGGCTAACCCGACC
ACCAACTCTTACAAACGTCTGGTTCCGGGTTACGAAGCTCCGGTT
ATGCTGGCTTACTCTGCTCGTAACCGTTCTGCTTCTATCCGTATCC
CGGTTGTTTCTTCTCCGAAAGCTCGTCGTATCGAAGTTCGTTTCCC
GGACCCGGCTGCTAACCCGTACCTGTGCTTCGCTGCTCTGCTGAT
GGCTGGTCTGGACGGTATCAAAAACAAAATCCACCCGGGTGAAGC
TATGGACAAAAACCTGTACGACCTGCCGCCGGAAGAAGCTAAAGA
AATCCCGCAGGTTGCTGGTTCTCTGGAAGAAGCTCTGAACGAACT
GGACCTGGACCGTGAATTCCTGAAAGCTGGTGGTGTTTTCACCGA
CGAAGCTATCGACGCTTACATCGCTCTGCGTCGTGAAGAAGACGA
CCGTGTTCGTATGACCCCGCACCCGGTTGAATTCGAACTGTACTA
CTCTGTTTAA

*argH*$^{m}$

ATGGCTCTGTGGGGTGGTCGTTTCACCCAGGCTGCTGACCAGCG
TTTCAAACAGTTCAACGACTCTCTGCGTTTCGACTACCGTCTGGCT
GAACAGGACATCGTTGGTTCTGTTGCTTGGTCTAAAGCTCTGGTTA
CCGTTGGTGTTCTGACCGCTGAAGAACAGGCTCAGCTGGAAGAA
GCTCTGAACGTTCTGCTGGAAGACGTTCGTGCTCGTCCGCAGCAG
ATCCTGGAATCTGACGCTGAAGACATCCACTCTTGGGTTGAAGGT
AAACTGATCGACAAAGTTGGTCAGCTGGGTAAAAAACTGCACACC
GGTCGTTCTCGTAACGACCAGGTTGCTACCGACCTGAAACTGTGG
TGCAAAGACACCGTTTCTGAACTGCTGACCGCTAACCGTCAGCTG
CAGTCTGCTCTGGTTGAAACCGCTCAGAACAACCAGGACGCTGTT
ATGCCGGGTTACACCCACCTGCAGCGTGCTCAGCCGGTTACCTTC
GCTCACTGGTGCCTGGCTTACGTTGAAATGCTGGCTCGTGACGAA

TCTCGTCTGCAGGACGCTCTGAAACGTCTGGACGTTTCTCCGCTG
GGTTGCGGTGCTCTGGCTGGTACCGCTTACGAAATCGACCGTGAA
CAGCTGGCTGGTTGGCTGGGTTTCGCTTCTGCTACCCGTAACTCT
CTGGACTCTGTTTCTGACCGTGACCACGTTCTGGAACTGCTGTCT
GCTGCTGCTATCGGTATGGTTCACCTGTCTCGTTTCGCTGAAGAC
CTGATCTTCTTCAACACCGGTGAAGCTGGTTTCGTTGAACTGTCTG
ACCGTGTTACCTCTGGTTCTTCTCTGATGCCGCAGAAAAAAAACCC
GGACGCTCTGGAACTGATCCGTGGTAAATGCGGTCGTGTTCAGG
GTGCTCTGACCGGTATGATGATGACCCTGAAAGGTCTGCCGCTGG
CTTACAACAAAGACATGCAGGAAGACAAAGAAGGTCTGTTCGACG
CTCTGGACACCTGGCTGGACTGCCTGCACATGGCTGCTCTGGTTC
TGGACGGTATCCAGGTTAAACGTCCGCGTTGCCAGGAAGCTGCTC
AGCAGGGTTACGCTAACGCTACCGAACTGGCTGACTACCTGGTTG
CTAAAGGTGTTCCGTTCCGTGAAGCTCACCACATCGTTGGTGAAG
CTGTTGTTGAAGCTATCCGTCAGGGTAAAGCTCTGGAAGACCTGC
CGCTGTCTGAACTGCAGAAATTCTCTCAGGTTATCGGTGAAGACG
TTTACCCGATCCTGTCTCTGCAGTCTTGCCTGGACAAACGTGCTG
CTAAAGGTGGTGTTTCTCCGCAGCAGGTTGCTCAGGCTATCGCTT
TCGCTCAGGCTCGTCTGGGTTAA

**Table S3.** pegRNA and gRNA sequences used in this study.

| pegRNA (gRNA) | Target | PAM | spacer | PBS | RTT |
|---|---|---|---|---|---|
| pegRNA01 | *glnA* | AGG | CTTGAACCTGG CACCCTGCA | AGGGTGC CAGGTT | GTCATAGCACTTGC |
| pegRNA02 | *glnA* | AGG | CTTGAACCTGG CACCCTGCA | AGGGTGC CAGGTT | GGTCATAGCTTGC |
| pegRNA03 | *glnA* | AGG | CTTGAACCTGG CACCCTGCA | AGGGTGC CAGGTT | GTCATAGCACTAC |
| pegRNA04 | *glnA* | TGG | TATTCGTTCTGA AATGTGTC | ACATTTCA GAACG | TCCATCACCTAGAC |
| pegRNA05 | *glnA* | TGG | TATTCGTTCTGA AATGTGTC | ACATTTCA GAACG | TTCCATCACAGAC |
| pegRNA06 | *glnA* | TGG | TATTCGTTCTGA AATGTGTC | ACATTTCA GAACG | TCCATCTACAGAC |
| pegRNA07 | *argH* | CGG | TACCGTTAGCG AGTTGCTGA | GCAACTC GCTAAC | CGGTTAGCATCGTG GCGTCA |
| gRNA-stop | *glnA* | TGG | GAGCAGGCACT GTACTACAT | - | - |
| gRNA-start | *glnA* | ATT | CCAGGAGAGTT AAAGTATGT | - | - |

**Table S4.** Strains used in this study.

| Name | Features [a] | Source |
|---|---|---|
| *E. coli* Nissle 1917 | Wild type strain (EcN). | From Ho lab |
| PR01 | EcN, DNA insertion at the locus 299 in *glnA*. | This study |
| PR02 | EcN, DNA deletion at the locus 299 in *glnA*. | This study |
| PR03 | EcN, DNA substitution at the loci 295, 297 and 298 in *glnA*. | This study |
| PR07 | EcN, DNA insertion at the locus 594 in *glnA*. | This study |
| PR08 | EcN, DNA deletion at the locus 594 in *glnA*. | This study |
| PR09 | EcN, DNA substitution at the loci 595 and 596 in *glnA*. | This study |
| PR10 | EcN, DNA insertion at the locus 391 in *argH*. | This study |
| PR13 | PR10, DNA insertion at the locus 299 in *glnA*. | This study |
| PR17 | EcN, early STOP codon insertion in *glnA*. | This study |
| PR18 | EcN, START codon exclusion in *glnA*. | This study |

[a.] Locus is counted from the first nucleotide of a specific gene.

**Table S5.** PR10/MG1655 abundance prediction by TIDER.

| PR10 /MG1655 (%) | PR10/MG1655 Abundance predicted by TIDER (%) | | | | Mean [b] | SD [c] |
|---|---|---|---|---|---|---|
| | Exp-1 [a] | Exp-2 | Exp-3 | Exp-4 | | |
| 0.0 | 0.0 | 0.0 | 0.0 | 0.0 | 0.0 | 0.0 |
| 1.0 | 0.1 | 0.1 | 0.1 | 0.1 | 0.1 | 0.0 |
| 5.0 | 0.5 | 0.3 | 1.2 | 0.3 | 0.6 | 0.4 |
| 10.0 | 9.1 | 6.9 | 9.1 | 7.9 | 8.3 | 0.9 |
| 20.0 | 8.6 | 9.8 | 9.9 | 7.6 | 9.0 | 0.9 |
| 50.0 | 51.0 | 48.5 | 50.9 | 52.9 | 50.8 | 1.6 |
| 100.0 | 100.0 | 100.0 | 100.0 | 100.0 | 100.0 | 0.0 |

[a.] Analysis of an individual sequencing result of one defined strain ratio.

[b.] The average of four abundance results.

[c.] Standard deviation of four abundance results.

**Table S6.** Primers used in this study.

| Primer | Sequence |
|---|---|
| **Primers for In-Fusion DNA assembly** | |
| XIA-PRC-107 | GACCGCCTACGGCGGCTGCGAAGGATCTAGGTGAAGATCC |
| XIA-PRC-108 | CGGGGAGGCAGACAAGGTATCCCTTTGATATGTAACGGTG |
| XIA-PRC-109 | GGATCTTCACCTAGATCCTTCGCAGCCGCCGTAGGCGGTC |
| XIA-PRC-110 | GCCGATGCTGTACTTCTTGTCCATCATGGTCTGTTTCCTGTGTG |
| XIA-PRC-111 | CACCGTTACATATCAAAGGGATACCTTGTCTGCCTCCCCG |
| XIA-PRC-112 | GACGGCAGCGAATTCGAGTGACCATGCGAGAGTAGGGAACT |
| XIA-PRC-113 | CATTTGAGAAGCACACGGTCACAGGAAACAGCTATGACCG |
| XIA-PRC-114 | TTGACTACCGGAAGCAGTGTTCTAGATTGTAAAACGACGGCCAGTC |
| XIA-PRC-115 | AGTTCCCTACTCTCGCATGGTCACTCGAATTCGCTGCCGTC |
| XIA-PRC-116 | CACACAGGAAACAGACCATGATGGACAAGAAGTACAGCATCGGC |
| XIA-PRC-132 | CACACAGGAAACAGACCATGATGGATAAGAAATACTCAATAGG |
| XIA-PRC-133 | TCCCAGCTGAGACAAATCAATGCGTGTTTCATAAAGACCAGTGAT |
| XIA-PRC-134 | TCTTTATGAAACACGCATTGATTTGTCTCAGCTGGGAGGTGACTC |
| XIA-PRC-135 | CCTATTGAGTATTTCTTATCCATCATGGTCTGTTTCCTGTGTG |
| XIA-PRC-136 | TGTCGATGCCATTGTTCCAC |
| XIA-PRC-137 | GTGGAACAATGGCATCGACA |
| XIA-PRC-142 | ACGGTTCTGAATTCGAATGACCATGCGAGAGTAGGGAACT |
| XIA-PRC-143 | AGTTCCCTACTCTCGCATGGTCATTCGAATTCAGAACCGT |
| XIA-PRC-160 | GTCACCTCCCAGCTGAGACAGGTCGATCCGTGTCTCGTAC |
| XIA-PRC-161 | GTACGAGACACGGATCGACCTGTCTCAGCTGGGAGGTGAC |
| XIA-PRC-162 | AATGCCGATATCCTATTGGCGGCTCTTGTATCTATCAGTG |
| XIA-PRC-163 | GTCTGACGCTCAGTGGAACG |

| | |
|---|---|
| XIA-PRC-164 | CCCTTAACGTGAGTTTTCGT |
| XIA-PRC-165 | GCAGGTATATGTGATGGGTTGAACATGTGAGCAAAAGGCC |
| XIA-PRC-166 | GGCCTTTTGCTCACATGTTCAACCCATCACATATACCTGC |
| XIA-PRC-167 | CACTGATAGATACAAGAGCCGCCAATAGGATATCGGCATT |
| XIA-PRC-168 | CACACAGGAAACAGACCATGATGGTGAGCAAGGGCGAGGAGGAT |
| XIA-PRC-169 | AGTTCCCTACTCTCGCATGGCTACTTGTACAGCTCGTCCA |
| XIA-PRC-170 | TGGACGAGCTGTACAAGTAGCCATGCGAGAGTAGGGAACT |
| XIA-PRC-171 | ATCCTCCTCGCCCTTGCTCACCATCATGGTCTGTTTCCTGTGTG |
| XIA-PRC-178 | CCTTGTCGCCTTGCGTATAACACGACGTTGTAAAACGACG |
| XIA-PRC-179 | CGTTGAATCGGGATATGCAGCACAGGAAACAGCTATGACC |
| XIA-PRC-180 | CGTCGTTTTACAACGTCGTGTTATACGCAAGGCGACAAGG |
| XIA-PRC-181 | GGTCATAGCTGTTTCCTGTGCTGCATATCCCGATTCAACG |
| XIA-PRC-194 | AACGACAGGAGCACGATCATCACGACGTTGTAAAACGACG |
| XIA-PRC-195 | CTGTTCGGTTCGTAAGCTGTCACAGGAAACAGCTATGACC |
| XIA-PRC-196 | GAGCATTGGCCACATGTGCTCCCTTAACGTGAGTTTTCGT |
| XIA-PRC-197 | CGTCGTTTTACAACGTCGTGATGATCGTGCTCCTGTCGTT |
| XIA-PRC-198 | GGTCATAGCTGTTTCCTGTGACAGCTTACGAACCGAACAG |
| XIA-PRC-199 | ACGAAAACTCACGTTAAGGGAGCACATGTGGCCAATGCTC |
| **Primers for colony PCR and sequencing** | |
| XIA-PRC-129 | GCGCTTCACCGATACCAAAG |
| XIA-PRC-130 | GAACACGTACTGACGATGCT |
| XIA-PRC-131 | TTCGCCCGGATGGATCTTGT |
| XIA-PRC-175 | ACGACTCACTGCGCTTTGAT |
| XIA-PRC-176 | CGATTTCATAGGCCGTTCCT |
| XIA-PRC-177 | CTGAACGTGCTGCTGGAAGA |

| | |
|---|---|
| XIA-PRC-186 | GCTCTGGTGATTCAGGACTC |
| XIA-PRC-187 | CAATGGTTCGTTCTCATGGC |
| XIA-PRC-200 | TGACAAAAGCGGTTCGGCAG |
| XIA-PRC-201 | TTCCAGTCGGGAAACCTGTC |
| XIA-PRC-202 | CCAGACTTTACGAAACACGG |
| XIA-PRC-203 | ACCGGTTTCGAATGCGTGGA |
| 113 | CATTTGAGAAGCACACGGTCACAGGAAACAGCTATGACCG |
| 114 | TTGACTACCGGAAGCAGTGTTCTAGATTGTAAAACGACGGCCAGTC |
| 253 | TGGTGTTGAAACGGGTAGCA |
| 254 | GCTACCCGTAACTCTCTGGA |
| M13F | GTAAAACGACGGCCAGT |
| M13R | GTCATAGCTGTTTCCTG |

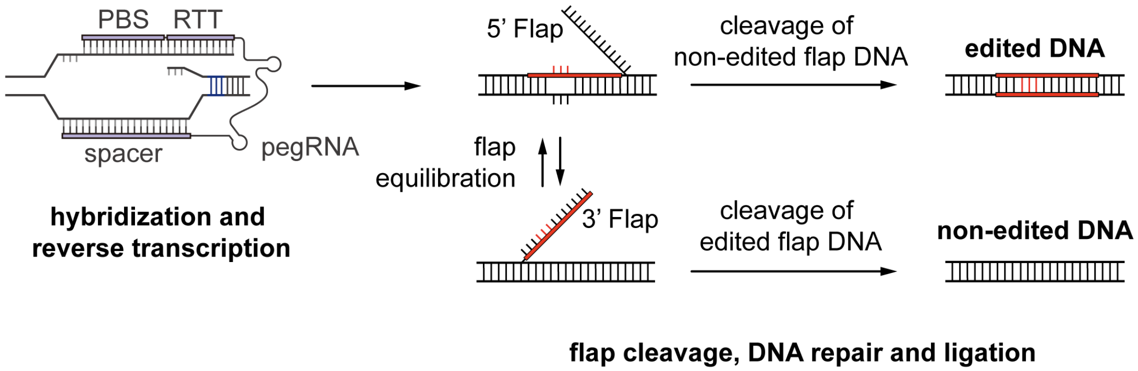


**Figure S1. Schematic illustration of prime editing mechanism.** PBS hybridizes to the exposed 3' end of the nicked DNA strand, and the reverse transcription of RTT is initiated to embed the designed edits in the 3' flap. The intended edits will be incorporated into the genome via DNA repair and 3' flap ligation when the 3' flap is retained and the 5' flap is cleaved during the flap equilibration process. PBS, primer binding site; RTT, reverse transcription template.

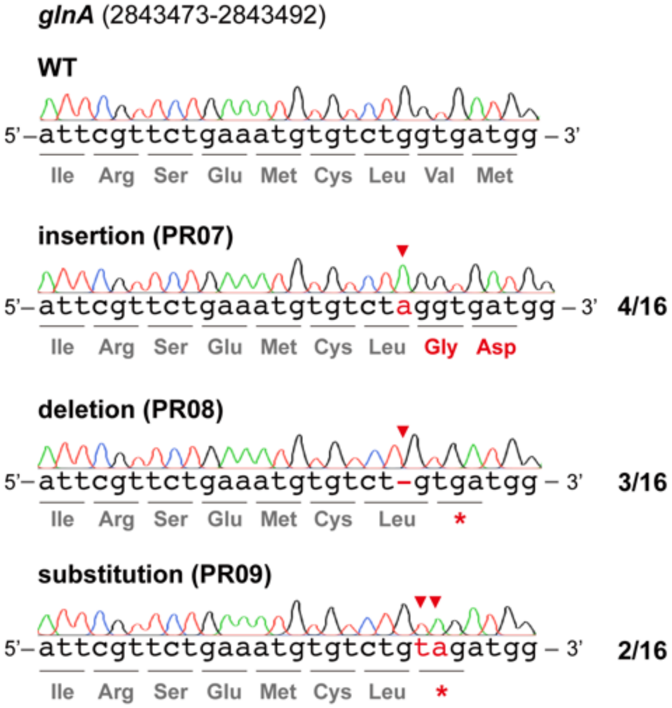


**Figure S2. Prime editing results at a different locus in *glnA*.** The editing was performed with pRC07, pRC08 and pRC09. The altered amino acids and the edited bases are highlighted in red, and the edited loci are indicated by the red arrows.

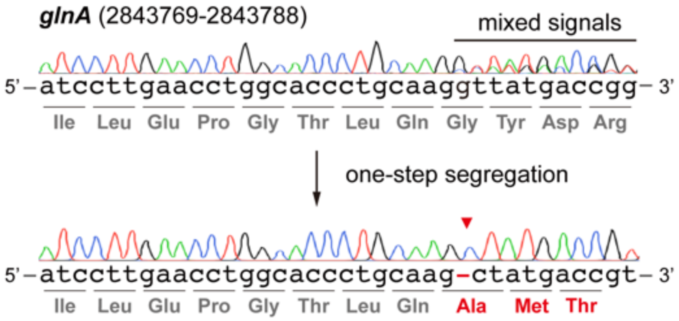


**Figure S3. One-step segregation for pure edited strains.** The altered amino acids and edited base are highlighted in red, and the edited position is indicated by the red arrow.

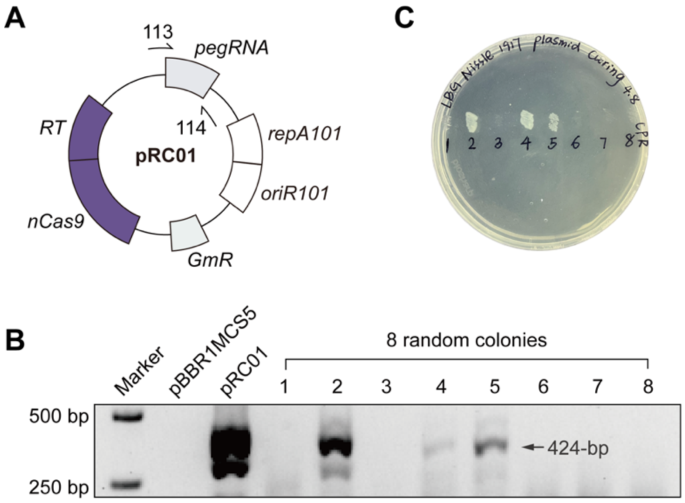


**Figure S4. Plasmid curing of EcN. (A)** Two primers, 113 and 114, were designed to amplify the specific sequences of pRC01 for verification. **(B)** Gel electrophoresis of plasmid cured and not cured colonies, with the backbone plasmid pBBR1MCS5 and pRC01 as negative and positive controls, respectively. **(C)** Sensitivity to gentamicin. The same colonies for gel electrophoresis were tested for gentamicin sensitivity. A colony with failed amplification and recovery sensitivity to gentamicin is considered plasmid cured.

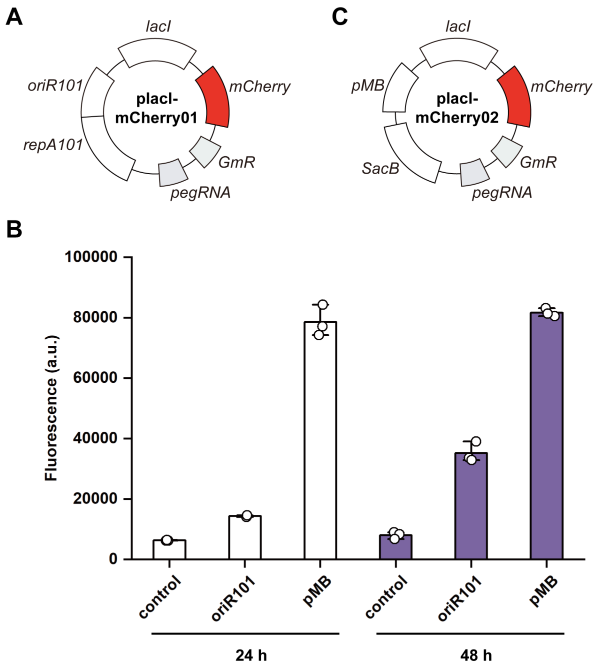


**Figure S5. Verification of the increased expression levels with extended induction time. (A)** The plasmid map of placI-mCherry01 and placI-mCherry02. **(B)** Fluorescence intensity. The fluorescence intensities for 24 h and 48 h induction are indicated in the white and purple columns, respectively. Three independent experiments were carried out and the standard deviations are represented by error bars.

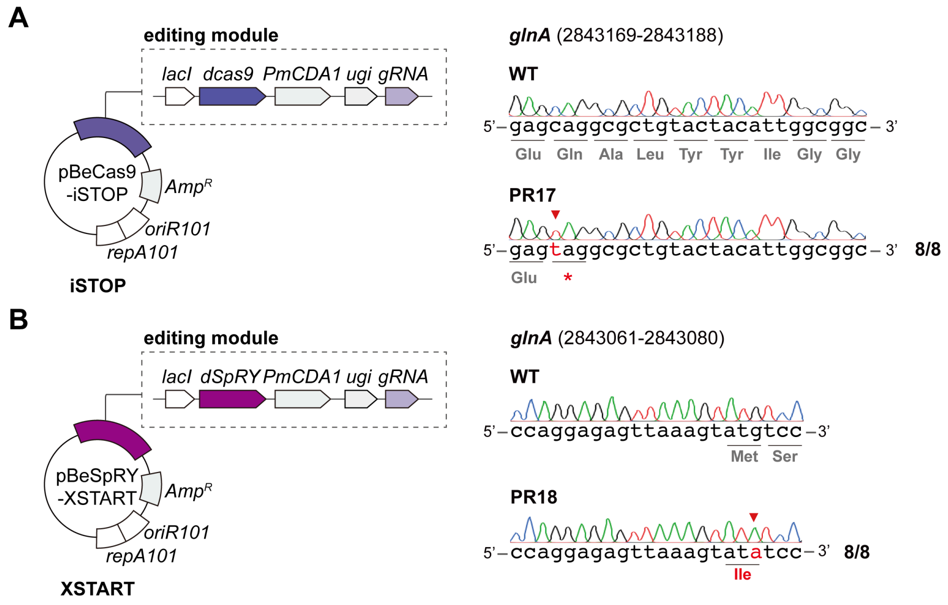


**Figure S6. Base editing system for iSTOP and XSTRAT. (A)** The plasmid map of pBeCas9-iSTOP and the sequencing result of PR17. The editing module contains *dCas9*, *PmCDA1* and *ugi*, driving by a *lacI*-$P_{trc}$ inducible system. The *gRNA* is controlled by $P_{J23119}$. **(B)** The plasmid map of pBeSpRY-XSTART and the sequencing result of PR18. The editing module contains *dSpRY*, *PmCDA1* and *ugi*, driving by a *lacI*-$P_{trc}$ inducible system. The *gRNA* is controlled by $P_{J23119}$. The altered amino acids and edited base are highlighted in red, and the edited position was indicated by the red arrow.

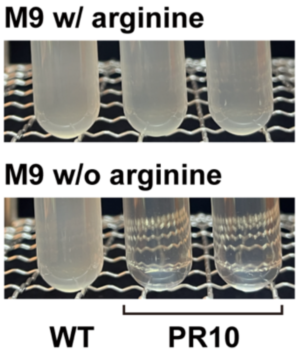


**Figure S7. Growth of the barcoded auxotrophic PR10 in M9 minimal medium with and without arginine.**

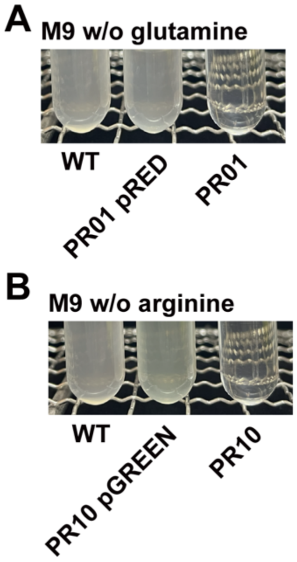


**Figure S8. Phenotypical evaluation of the ARG-free platform. (A)** Growth of PR01 with and without pRED in M9 medium without glutamine, and **(B)** growth of PR10 with and without pGREEN in M9 medium without arginine.

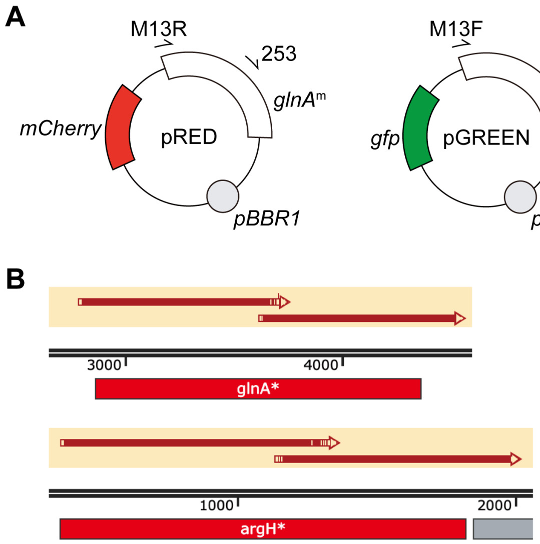


**Figure S9. Design and construction of pRED and pGREEN. (A)** Illustration of the two plasmids with sequencing primers highlighted on pRED (M13R and 253) and pGREEN (M13F and 254). **(C)** Sequencing verification of *glnA*$^m$ and *argH*$^m$ in pRED and pGREEN.

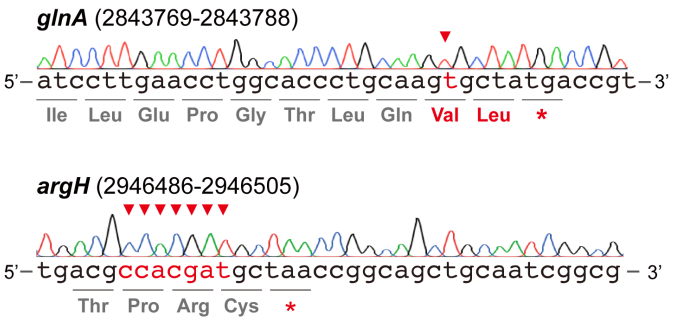


**Figure S10. Sequencing results of the double-auxotrophic PR13.** The altered amino acid and the edited base are highlighted in red, and the edited positions are indicated by red arrows.

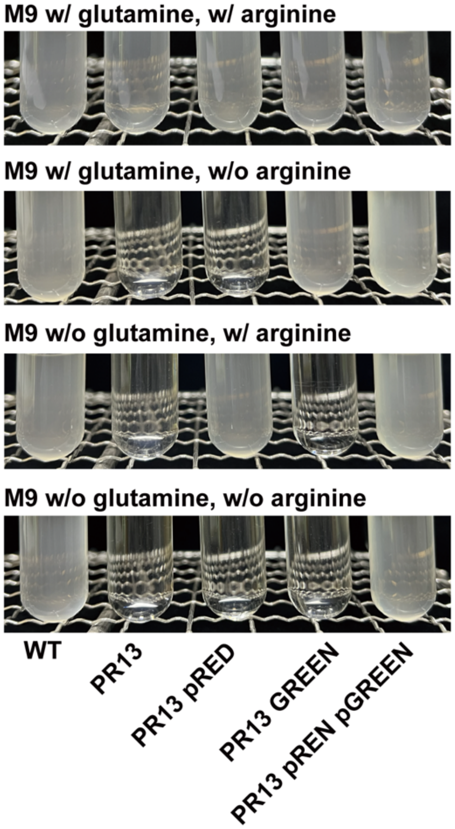


**Figure S11. Phenotypical evaluation of the double-auxotrophic PR13 with pRED or/and pGREEN in M9 medium with and without glutamine and arginine.**

### Reference


1. K. A. Datsenko, B. L. Wanner, One-step inactivation of chromosomal genes in *Escherichia coli* K-12 using PCR products. *Proc. Natl. Acad. Sci. U. S. A.* **97**, 6640-6645 (2000).
2. Y. Wei, L. J. Feng, X. Z. Yuan, S. G. Wang, P. F. Xia, Developing a base editing system for marine *Roseobacter* clade bacteria. *ACS Synth. Biol.* **12**, 2178-2186 (2023).
3. Y. Chen *et al.*, Self-replicating shuttle vectors based on pANS, a small endogenous plasmid of the unicellular cyanobacterium *Synechococcus elongatus* PCC 7942. *Microbiology* **162**, 2029-2041 (2016).
4. A. V. Anzalone *et al.*, Search-and-replace genome editing without double-strand breaks or donor DNA. *Nature* **576**, 149-157 (2019).
5. X. Li, Y. Wei, S. Y. Wang, S. G. Wang, P. F. Xia, One-for-all gene inactivation via PAM-independent base editing in bacteria. *bioRxiv* [preprint] (2024). https://doi.org/10.1101/2024.06.17.599441 (accessed 15 October 2024).
